# Tracking the hidden dynamics of proprioception

**DOI:** 10.64898/2026.08.16.745118

**Authors:** Pamela Villavicencio, Dominik Straub, Mabel Ziman, Matthias Will, Roberta Klatzky, Cristina de la Malla, Jonathan S. Tsay

## Abstract

Every movement unfolds with a simple question: Where is my body? The nervous system answers through proprioception—the sense of limb position (static position sense) and movement (dynamic proprioception). Although position sense has been well characterized, dynamic proprioception has remained difficult to isolate and measure. Here we introduce a continuous proprioceptive tracking paradigm, coupled with computational modelling, that captures dynamic proprioception in real time. We first establish that this approach is sensitive, reliable and efficient. Leveraging this method, we then show that dynamic proprioception provides faster and more faithful estimates of limb state than vision, dominates multisensory state estimation when vision is also available, and is not correlated with conventional measures of position sense. Together, these findings provide a new quantitative framework for characterizing dynamic proprioception in health and disease.

## Introduction

*“No science attains maturity until it acquires systems of measurement.”*

Logan Clendening

Imagine trying to button a shirt, type on a keyboard, or take a step without knowing where your body is. Even the simplest actions would become extraordinarily difficult. This sense of the body—proprioception—tells us where our limbs are (static position sense) and how they are moving (dynamic proprioception). When proprioception is impaired, movements become dramatically slower, less accurate, and more effortful (Ghez et al., 1995; Proske & Gandevia, 2012; Rothwell et al., 1982; Sainburg et al., 1993, 1995; Sanes et al., 1985; Sarlegna et al., 2006; Schaffer et al., 2021). Every movement therefore depends on the nervous system continuously tracking the body’s posture.

Yet most of what we know about proprioception comes from empirical studies that provide static snapshots of stationary posture. Conventional assays ask participants to report where their limbs are at a specific point in time, providing discrete measures of position sense (Block & Liu, 2023; Clayton et al., 2014; Kuling et al., 2013, 2016; Mostafa et al., 2019; Tsay et al., 2021, 2024). Approaches that probe dynamic proprioception typically ask participants to reproduce previously experienced movements or initiate an action when they perceive their body to have reached a specific position, introducing confounding contributions from working memory, motor noise, feedforward efferent predictions, and effort (Bhanpuri et al., 2013; Callaghan & Reinkensmeyer, 2025; Cordo, 1990; Goble & Brown, 2009; Ohashi et al., 2019; Tulimieri & Semrau, 2023; A. L. Wong et al., 2024; Zangakis et al., 2026). Thus, dynamic proprioception—how the nervous system continuously estimates the body’s changing configuration during ongoing movement— has remained difficult to isolate and measure.

To overcome these limitations, we developed a new continuous proprioceptive tracking task that measures how accurately and quickly the nervous system tracks the body during movement. A robotic device passively moved one arm along an unpredictable path, generating continuously changing proprioceptive signals that participants attempted to track with the other arm. The correspondence between the two arms provided a moment-to-moment behavioral proxy of dynamic proprioception, with minimal demands on working memory due to continuous, real-time tracking and minimal contribution from efferent prediction due to the passive, unpredictable movement of the tracked hand.

Yet behavioral tracking performance does not reflect proprioception alone: expressing these dynamic state estimates requires active movement of the tracking hand, introducing variability from motor execution and effort-related costs. We therefore paired the task with computational modeling as an initial approach to dissociating proprioceptive uncertainty from these motor contributions (Straub & Rothkopf, 2022). We first establish that this approach provides a sensitive, reliable, and efficient measure of dynamic proprioception, and then use it to reveal several defining properties of proprioception during ongoing movement. Together, these findings provide a new quantitative framework for characterizing dynamic proprioception in health and disease.

## Results

### Proprioceptive tracking provides a sensitive assay of dynamic proprioception

We developed a continuous proprioceptive tracking paradigm that provides a behavioral assay of dynamic proprioception—the rapid, continuous, and largely automatic estimation of body state during movement. Across two sessions, participants used their left hand to continuously track the passive, unpredictable displacement of their right hand, which served as the proprioceptive target (Figure 1a; Experiment 1, N = 19). As illustrated by a representative trial (Figure 1b), movements of the active left hand closely followed those of the passively displaced right hand.

**Figure 1.**
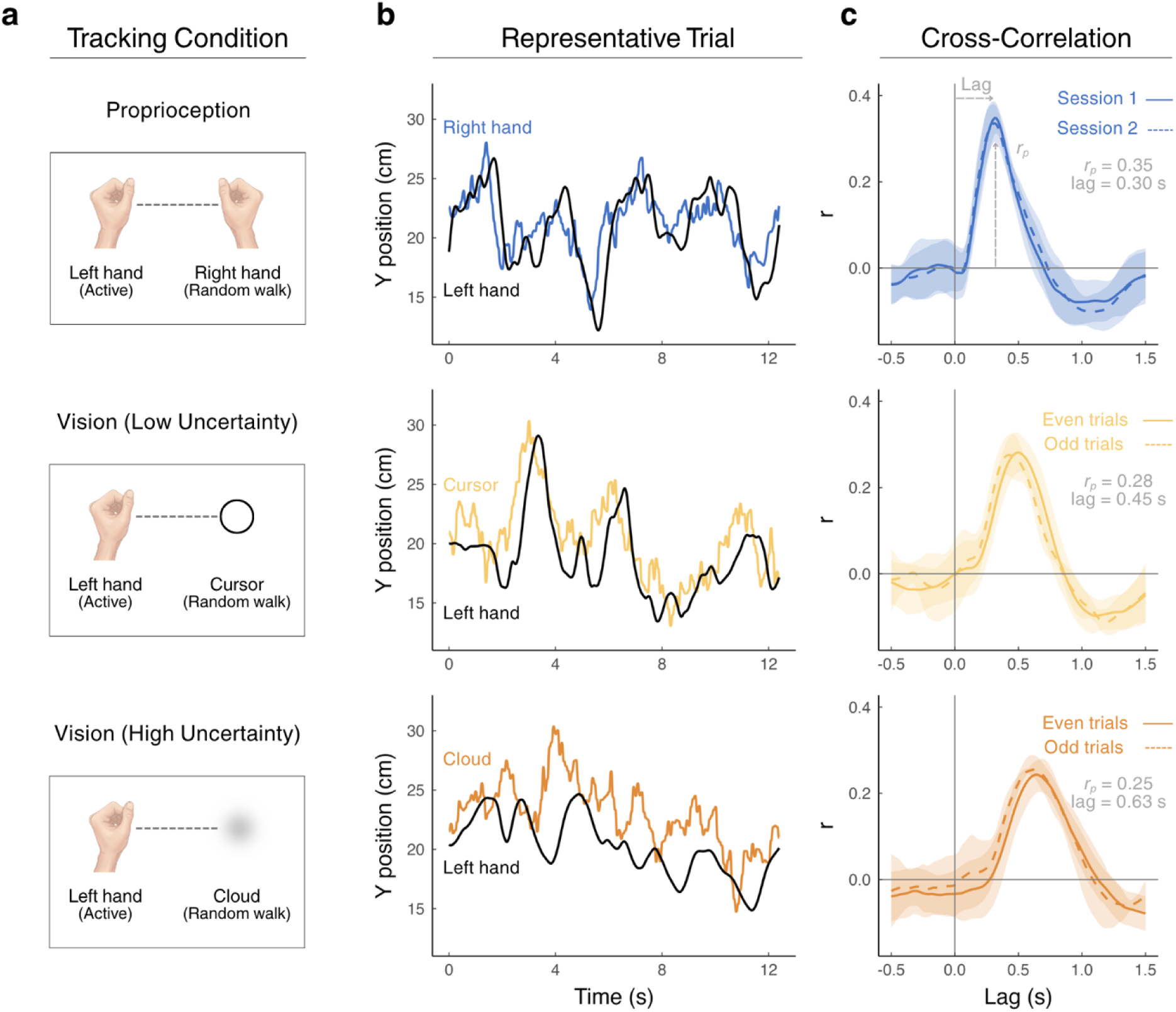
Proprioceptive tracking provides a sensitive assay of dynamic proprioception. **(a)** Participants used their left hand to actively track a proprioceptive target (right hand) or a visual target (cursor or cloud). In both conditions, the target moved along the forward-backward y-axis according to a random-walk trajectory (range along the y-axis: 12–35 cm; with smaller values corresponding to positions closer to the body). Both hands were occluded from view. **(b)** Representative tracking behavior from a single participant. Colored traces indicate the target trajectory (blue, proprioceptive target; yellow, visual cursor target; orange, visual cloud target), whereas black traces show the movements of the active left hand as it tracked the corresponding target. **(c)** Cross-correlations between the target and tracking velocities for the representative participant. For proprioceptive tracking, cross-correlations are shown separately for the two testing sessions on different days. For visual tracking, cross-correlations are shown separately for even-and odd-numbered trials within a single session. Solid lines and shaded ribbons denote the mean ± SD. The average peak correlation (*r_p_*) and lag for each tracking condition are reported for this participant.

To quantify tracking performance, we computed the cross-correlation between the velocity of an invisible proprioceptive target (right hand) and the velocity of an invisible active left-hand tracking response (Figure 1c). From the cross-correlation function for each trial, we extracted two metrics: peak correlation (*r_p_*), reflecting the fidelity of dynamic proprioception; and lag, reflecting the latency with which proprioception guided movement (Burge & Bonnen, 2025). For the representative participant, tracking was both robust (mean ± SD: *r_p_* = 0.36 ± 0.04) and rapid (lag = 0.30 ± 0.03 s).

These features were consistent across participants (Figure 2a), with all participants exhibiting strong target–tracking coupling (*r_p_*, mean ± SEM = 0.35 ± 0.08) and relatively short response delays (lag = 0.27 ± 0.04 s). Both metrics exhibited marked between-subject variability without evidence of floor or ceiling effects, underscoring the paradigm’s sensitivity to meaningful individual differences. Notably, the mean response delay of 270 ms falls squarely within the temporal range of rapid motor control (Flanders & Cordo, 1989; Scott, 2016), demonstrating that proprioceptive tracking sensitively captures the dynamics required to guide ongoing movement.

**Figure 2.**
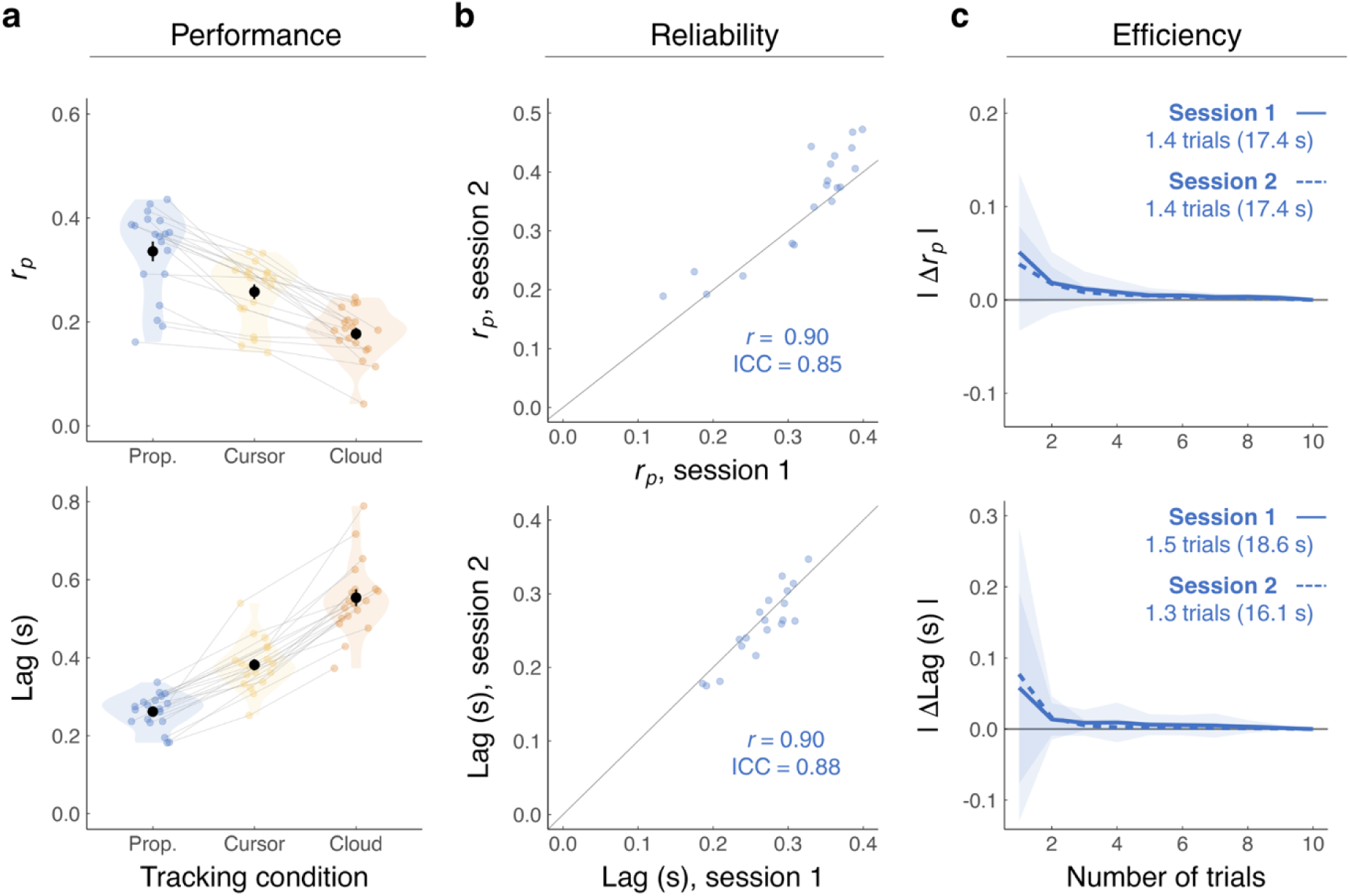
Proprioceptive tracking provides reliable and efficient estimates of dynamic proprioception. **(a)** Cross-correlation metrics for proprioceptive and visual tracking conditions. Colored points denote individual means across different tracking conditions; black points and error bars denote group means ± SEM. **(b)** Test–retest reliability of the proprioceptive tracking condition across two testing sessions separated by over one week. Grey lines show the identity line. Points denote individual means. Between-session Pearson’s r and ICC values are reported. **(c)** Absolute deviation of peak correlation and lag from the reference estimate obtained using all proprioceptive trials, plotted as a function of the number of trials included in the bootstrap resampling. Solid lines and shaded ribbons indicate bootstrap means ± SD. The number of trials and corresponding testing time (12.4 s per trial) required for the absolute deviation to fall below 5% of the reference estimate are reported.

We next examined additional features of tracking performance. Across participants, the mean positional difference between the tracking and target hands was small, with a slight tendency to undershoot the target (Session 1: −0.6 ± 0.3 cm; t-test vs. 0: *t*(18)=-2.4, *p* = 0.03, *D* =-0.6; Session 2: −0.5 ± 0.3 cm; t-test vs. 0: *t*(18)=-1.6, *p* = 0.1). This modest undershoot may reflect effort-related costs, whereby the cost of larger movements favors responses that do not fully reproduce the extent of the target trajectory. RMSE (root mean squared error) remained stable from early to late trials (*F*(1, 354) = 1.2, *p* = 0.269), providing little evidence of fatigue despite sustained tracking demands. Finally, active tracking trajectories contained fewer high-frequency components than the passive right-hand movement, consistent with a low-pass transformation (*t*(18) =-3.1, *p* = 0.003, *D* = - 0.7). This attenuation of rapid fluctuations may reflect both biomechanical constraints on the tracking arm and temporal filtering of noisy sensory estimates.

These features highlight an important distinction: continuous tracking performance provides a behavioral proxy of dynamic proprioception, but not a direct measure of the latent proprioceptive estimate. The observed trajectory reflects not only sensory uncertainty, but also biomechanical constraints, control costs, and motor execution noise involved in transforming sensory input into action. We therefore use computational modeling in a later section to account for these downstream influences and recover the latent sensory uncertainty associated with dynamic proprioception.

### Proprioceptive tracking provides reliable estimates of dynamic proprioception

A critical requirement for any new behavioral construct is psychometric reliability. This has proven challenging for many existing proprioceptive assays, which often exhibit only modest reliability, limiting their utility for studying individual differences, tracking longitudinal change, and quantifying impairment in clinical populations (Carey et al., 1996; Elangovan et al., 2014; Fatoye et al., 2008; Horváth et al., 2023; Juul-Kristensen et al., 2008; Lönn et al., 2000). We therefore asked whether proprioceptive tracking yields reliable estimates of dynamic proprioception.

To answer this question, we evaluated the test–retest reliability of our proprioceptive metrics across two testing sessions separated by over one week (mean ± SD: 8 ± 3 days). For our representative participant, strikingly, cross-correlation functions were identical across sessions (Figure 1c; Session 1: *r_p_* = 0.36 ± 0.03, lag = 0.30 ± 0.03 s; Session 2: *r_p_* = 0.36 ± 0.04, lag = 0.30 ± 0.03 s). This consistency generalized across participants, with estimates of peak correlation and response lag clustering tightly around the identity line (Figure 2b; *r_p_*: ICC(A,1) = 0.85, 95% CI [0.48–0.95]; lag: ICC(A,1) = 0.88, 95% CI [0.71–0.95]). Together, these findings establish proprioceptive tracking as a highly reliable assay of dynamic proprioception.

### Proprioceptive tracking provides efficient estimates of dynamic proprioception

Many existing proprioceptive assessments require a large number of trials to obtain stable estimates (Cressman et al., 2021; Elangovan et al., 2014; Lowrey et al., 2020). Concretely, conventional assessments of static position sense often require more than 400 trials, corresponding to 15–60 minutes of testing (van Beers et al., 1998). These demands limit their utility as rapid screening tools, constrain studies requiring repeated measurements, and hinder large-scale investigations (Valero-Cuevas et al., 2024). An ideal assay would provide stable proprioceptive estimates with minimal testing time. We therefore asked how much data are required to obtain stable behavioral estimates of dynamic proprioception.

To answer this question, we performed a bootstrap convergence analysis of both cross-correlation metrics to determine the minimum amount of data required to obtain stable estimates. For each participant and session, cross-correlation metrics were estimated using increasing number of trials and compared with reference values derived from all available trials (10 trials/session; ∼10 min of testing). Convergence was defined as the smallest number of trials for which the metric’s deviation fell below 5% of the reference value. Strikingly, stable estimates of peak correlation and lag were obtained after only 1.4 ± 0.9 trials, and 1.4 ± 1.2 trials (mean ± SD), respectively, corresponding to less than 20 seconds of testing (Figure 2c). We found that 18/19 participants reached convergence on both metrics within 3 trials (less than 40 seconds of testing). Together, these findings establish continuous proprioceptive tracking as a highly efficient assay of dynamic proprioception, whereby stable estimates can be obtained from an assessment lasting less than a minute.

### Dynamic proprioception guides movement faster and more precisely than vision

Armed with this new behavioral assay, we next sought to define the properties of dynamic proprioception. In tasks assaying position sense, vision is often regarded as the dominant modality, exerting a stronger influence on state estimation than proprioception (Gaffin-Cahn et al., 2019; Hagura et al., 2007; Welch & Warren, 1986) (but see: Mon-Williams et al., 1997; Plooy et al., 1998; van Beers et al., 2002). Yet natural behavior is continuous, rapid, and automatic. Under these conditions, the contribution of each sensory modality depends not only on its precision but also on how rapidly it can support ongoing movement (Crevecoeur et al., 2016; A. L. Wong et al., 2024). Whether vision or dynamic proprioception provides the faster and more precise estimate of body state during movement therefore remains an open question.

To answer this question, we used our tracking paradigm to directly compare the speed and precision of proprioceptive and visual state estimation during movement. In addition to the proprioceptive tracking condition, participants completed two visual tracking conditions in which they tracked either a low-uncertainty visual cursor or a high-uncertainty visual cloud (Figure 1a). Consistent with prior work (Ambrosi et al., 2022; Bonnen et al., 2015; Burge & Bonnen, 2025; Burge & Cormack, 2024), continuous tracking was sensitive to graded differences in visual uncertainty. Specifically, tracking the cursor produced higher peak correlations (*t(*72) = 5.9, *p* < 0.0001, *D_z_* = 1.8) and shorter lags (*t(*72) = −12.5, *p* < 0.0001, *D_z_* = −2.8) compared to tracking the cloud. On average, tracking the cursor was 31% faster and 46% more precise than tracking the cloud (cursor *r_p_* = 0.26 ± 0.01, lag = 0.38 ± 0.01 s; cloud *r_p_* = 0.18 ± 0.01, lag = 0.55 ± 0.02 s).

Strikingly, dynamic proprioception supported faster and more precise tracking than vision. Relative to both visual targets, proprioceptive tracking yielded higher peak correlations (vs. cursor: *t*(72) = 5.7, *p* < 0.0001, *D_z_* = 1.7; vs. cloud: *t*(72) = 11.6, *p* < 0.0001, *D_z_* = 2.4), and shorter response lags (vs. cursor: *t*(72) =-8.7, *p* < 0.0001, *D_z_* =-2.6; vs. cloud: *t*(72) =-21.2, *p* < 0.0001, *D_z_* =-3.5). On average, proprioceptive tracking was 31% faster than tracking the visual cursor and 53% faster than tracking the visual cloud. This advantage in dynamic proprioception was remarkably consistent, with all 19 participants exhibiting the same pattern (Figure 1c & Figure 2a). Together, these findings indicate that dynamic proprioception furnishes the nervous system with a faster and more faithful estimate of body state during naturalistic movement.

### Dynamic proprioception dominates multisensory state estimation during movement

The previous experiments establish not only that dynamic proprioception provides continuous estimates of body state, as evidenced by accurate, short-latency tracking, but also that vision provides substantially slower and less precise estimates. This raises a fundamental question: What does vision contribute when dynamic proprioception is also available?

We considered four possibilities. A **proprioception-only** account predicts that state estimates rely exclusively on proprioception, with vision effectively discarded when proprioceptive information is available (Crevecoeur et al., 2016; A. L. Wong et al., 2024). Such dominance can emerge when sensory modalities differ markedly in precision, such that perception is governed by the more reliable signal.

Second, a **reliability-weighted integration** account predicts that proprioceptive and visual signals are combined according to their relative reliability: the more precise signal receives greater weight, while the noisier signal contributes less (Smeets et al., 2006; van Beers et al., 1999, 2002) (also see: Alais & Burr, 2004; Bays & Wolpert, 2007; Ernst & Banks, 2002; Ernst C Bülthoff, 2004). Under this account, proprioception should exert greater influence than vision because it is more precise, but vision should nevertheless make a measurable contribution to the resulting state estimate.

Third, an **equal-weighting** account predicts that proprioceptive and visual signals are combined using a simple 50/50 heuristic, with each contributing equally regardless of its reliability (van Atteveldt et al., 2014). Such a strategy would produce multisensory estimates intermediate between the two unimodal signals, even when their precision differs substantially.

Finally, a **visual-only** account predicts that multisensory state estimates are governed only by vision, with proprioception contributing little when visual information is available. Although this possibility runs counter to the superior precision and shorter latency of dynamic proprioception observed here, visual capture has been reported in other multisensory contexts and therefore provides an important competing account (Posner et al., 1976; Rock & Victor, 1964).

Our continuous tracking paradigm provides a test of these predictions. Participants tracked a multisensory target in which visual (cursor or cloud, tested in separate sessions) and proprioceptive targets were presented simultaneously (Figure 3a). During the aligned phase, the two targets occupied the same location. During the offset phase, a visuo-proprioceptive conflict was gradually introduced by shifting the visual target either 5 cm ahead of or behind the proprioceptive target (direction counterbalanced across participants). No participant reported noticing the offset. We quantified multisensory tracking using the cross-correlation metrics (*rₚ* and lag) and mean tracking position relative to the proprioceptive target, with the position of the right hand defined as 0 cm.

**Figure 3.**
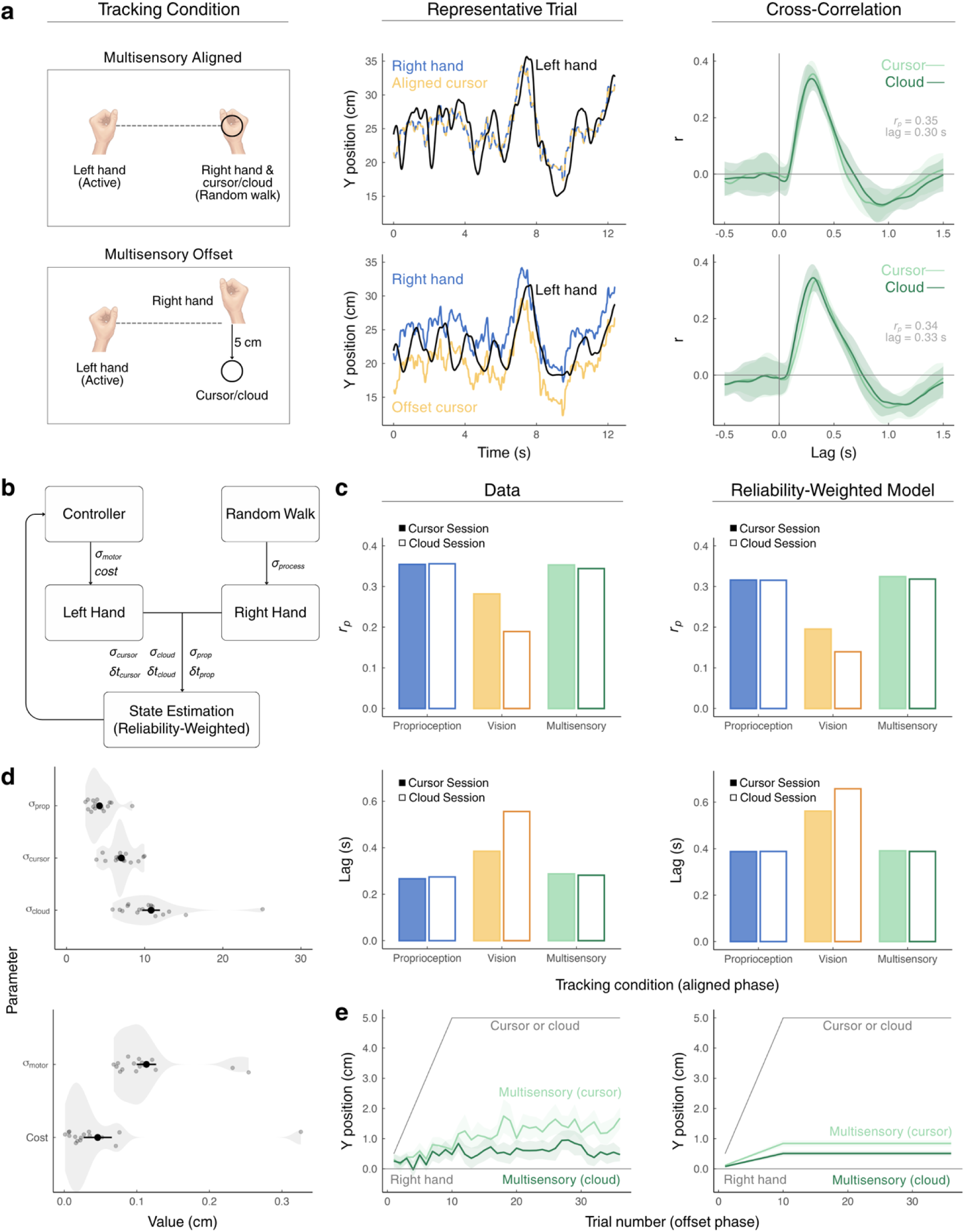
Dynamic proprioception dominates multisensory state estimation during movement. **(a)** Multisensory tracking in the aligned and offset phases. Participants used their left hand to track their passively moved right hand (a proprioceptive target) and a simultaneously presented visual target (cursor or cloud, tested on separate days). When the proprioceptive (blue) and visual (yellow) targets were aligned, the representative participant tracked the right hand closely; under an inter-sensory conflict (offset phase) — in which the visual target was displaced 5 cm forward or backward relative to the right hand — responses settled between the two targets, biased toward the right hand. This participant showed cross-correlations similar to those seen with proprioceptive tracking (lines and ribbons denote mean ± SD); mean *rₚ* and lag are reported. **(b)** Schematic of the reliability-weighted model. During continuous tracking, the discrepancy between the target and the actively moved tracking hand is observed through noisy proprioceptive (*σ_prop_*) and visual (*σ*_cursor_ or *σ*_cloud_) signals subject to sensory delays ($), fixed at 75 ms for proprioception and 150 ms for vision. These signals are combined via reliability-weighting yielding a single integrated estimate dominated by the more precise (proprioceptive) modality. A controller then generates motor commands to minimize the discrepancy between this estimate and the target, subject to control costs and motor noise (*σ_m_*_otor_). **(c)** Median *r_p_* (top) and lag (bottom) values during the aligned phase, for the data (left) and for model simulations generated from best-fit parameters (right). Filled and open bars denote sessions in which the visual target was the cursor and the cloud, respectively. **(d)** Posterior mean estimate of each model parameter for each participant (translucent points), and across participants (black points and error bars, mean ± SEM). **(e)** Multisensory tracking during the offset phase, in the data (left) and model simulations (right). The average y-position of the left (tracking) hand is shown relative to the proprioceptive right hand (horizontal grey line, y = 0 cm). The visual target was gradually displaced 5 cm from the right hand (direction counterbalanced across participants), as indicated by the rising grey line. Colored lines and ribbons denote mean ± SEM.

The aligned phase revealed clear proprioceptive dominance. Multisensory tracking was statistically indistinguishable from proprioceptive tracking alone, whether the visual target was a cursor (*r_p_*: *t*(72) =-0.6, *p* = 0.98; lag: *t*(72) =-1.2, *p* = 0.73) or a cloud (*r_p_*: *t*(72) = 0.3, *p* = 1.0; lag: *t*(72) =-0.3, *p* = 1.0). Multisensory tracking was, however, substantially more accurate and faster than either visual condition (vs. cursor *r_p_*: *t*(72) =-6.3, *p* <.0001, *D*_z_ =-1.5; lag: *t*(72) = 7.5, *p* <.0001, *D*_z_ = 2.0; vs. cloud *r_p_*: *t*(72) =-11.3, *p* <.0001, *D*_z_ =-2.0; lag: *t*(72) = 21.0, *p* <.0001, *D*_z_ = 3.2). Thus, multisensory state estimation closely resembled proprioception alone, arguing against the vision-only and equal-weighting accounts. However, because reliability weighting would also strongly favor the more precise proprioceptive signal, the aligned condition could not sensitively distinguish reliability weighting from exclusive reliance on proprioception.

The offset phase confirmed that proprioception remained dominant while revealing that a small contribution from vision emerged in the case of cue conflict. If state estimation relied exclusively on proprioception, shifting the visual target should have had no effect. Instead, tracking shifted modestly toward vision while remaining much closer to the proprioceptive target (Figure 3e). Averaged across both multisensory conditions, tracking position was 1.0 ± 0.2 cm (mean ± SEM), lying between the proprioceptive (0 cm) and visual (5 cm) targets but significantly below the 2.5-cm midpoint (*t*(18) =-7.6, *p* < 0.001, *D* =-1.7). Thus, proprioception strongly dominated the multisensory estimate, although vision was not entirely discarded under cue conflict.

Moreover, consistent with reliability-weighted integration, this smaller visual contribution depended on its precision. Participants shifted more than twice as far toward the low-uncertainty cursor (1.3 ± 0.3 cm; *t*(18) = 4.1, *p* = 0.0007, *D* = 0.9) than the high-uncertainty cloud (0.6 ± 0.2 cm; *t*(18) = 2.4, *p* = 0.025, *D* = 0.6) (Figure 3e). Trial-by-trial analyses further suggested that this divergence emerged late in the offset phase, with significant cursor–cloud differences observed on several trials between trials 23 and 36 (ts <-2.23, ps < 0.038). Together, these behavioral results establish that dynamic proprioception dominates multisensory state estimation, while vision exerts a smaller influence that varies with its precision.

We next asked whether this behavioral pattern could be quantitatively captured by the different models of continuous tracking (Figure 3b), extending the framework of Straub and Rothkopf (2022). Specifically, the behavioral results indicate that the proprioceptive dominance model should be favored under the aligned condition, but given cue conflict, the model should reveal a small weight for visual input consistent with its precision. Note that it is not to be expected that the same model would apply to both conditions, as spatial alignment has been shown to be critical to optimally weighted integration of vision and touch (Gepshtein et al., 2005).

All models assume that the observer receives delayed, noisy observations of both the target (visual, proprioceptive, or both) and the active tracking hand (proprioceptive). The target evolves according to a random walk with fixed process noise (*σ_process_*), whereas each sensory modality is corrupted by observation noise (*σ_prop_*, *σ_cursor_*, *σ_cloud_*). A Kalman filter combines sensory observations with the predicted target state. The models differed only in how visual and proprioceptive signals contributed to this estimate: the proprioception-only model relied exclusively on proprioception, the vision-only model exclusively on vision, the equal-weighting model assigned equal weight to both signals, and the reliability-weighting model weighted them according to their relative precision. In all models, a controller generates motor commands that minimize tracking error while penalizing movement effort (control cost, *c*), with commands corrupted by execution noise (*σ_motor_*). The models additionally incorporate realistic biomechanics of the tracking arm using point-mass dynamics that capture inertia, damping, and muscle-force dynamics (Kasuga et al., 2022).

We fit each model separately to each participant’s aligned-phase data and evaluated its ability to account for behavior in both the aligned and offset phases. For clarity, we feature the reliability-weighting model in Figure 3; results for all four models are reported in Figure S1 and Table S1. Qualitatively, reliability weighting was the only model to reproduce the key behavioral signatures across both phases: strong proprioceptive dominance when the cues were aligned, a modest visual influence under cue conflict, and a larger visual influence when vision was more precise.

Quantitative model comparison provided a more nuanced picture. Vision-only and equal-weighting models were clearly disfavored in both the aligned (ΔELPD-LOO, mean ± SEM: vision-only =-349 ± 91; equal weighting =-131 ± 26) and offset phases (ΔELPD: vision-only =-24598 ± 2731; equal weighting =-9115 ± 1489). Among the remaining models, the aligned phase favored proprioception-only over reliability weighting (ΔELPD-LOO: proprioception-only = 62 ± 13). In the offset phase, however, model comparison did not clearly distinguish proprioception-only from reliability weighting (ΔELPD: proprioception-only = 1323 ± 756), despite the behavioral evidence for a small, precision-dependent visual contribution.

We recognize this apparent discrepancy between the behavioral and modeling results. Rather than undermining the behavioral findings, this discrepancy helps define where the current models can—and cannot yet—account for the observed dynamics. The model comparison robustly rules out vision-only and equal-weighting accounts and confirms proprioception-only when cues align, but is less decisive between proprioception-only and reliability weighting under cue conflict conditions. This ambiguity likely reflects remaining model limitations: none of the candidate models accurately reproduced the tracking dynamics (Figure S1). Thus, further refinement will be needed to more precisely resolve the integration scheme underlying proprioceptive dominance, a point we return to in the Discussion. Regardless of this mechanistic distinction, the behavioral and computational results converge on a central conclusion: dynamic proprioception dominates multisensory state estimation during movement.

### Computational modeling isolates latent dynamic proprioceptive estimates from downstream motor and cognitive processes

A longstanding limitation of conventional assessments is that estimates of proprioceptive function are often confounded by motor execution noise, working memory, and effort-related processes (Callaghan & Reinkensmeyer, 2025; Goble & Brown, 2009; Ohashi et al., 2019; Tulimieri & Semrau, 2023; Wong et al., 2024; Zangakis et al., 2026). For example, in retrospective trajectory reproduction tasks, performance reflects not only the precision of proprioception, but also memory of the trajectory, motor execution noise, and the effort invested in reproducing it accurately.

Importantly, our behavioral measures are not immune to these same confounds. Peak cross-correlation, response lag, and tracking position all reflect the combined influence of sensory, motor, and control processes. Working-memory demands, however, are minimized by the continuous nature of the task, as proprioceptive estimates are used online rather than retained for later report. This is where the computational modeling introduced earlier provides a critical advantage: it allows us to move beyond the observed behavior to infer the latent sensory estimate that gave rise to it. By specifying a normative model of continuous tracking that explicitly captures distinct sources of variability, we can recover latent proprioceptive uncertainty and dissociate it from motor variability, control costs, and biomechanical constraints.

Figure 3d shows the posterior mean of each parameter, estimated per participant from the reliability-weighted model (comparable estimates were obtained under all four models; Table S1). Consistent with our behavioral analyses, proprioceptive uncertainty was lowest (mean ± SEM: 4.2 ± 0.4 cm), followed by uncertainty for the visual cursor (7.0 ± 0.4 cm) and then the visual cloud (10.8 ± 1.1 cm) (*F*(2,30) = 22.9, *p* < 0.0001, *η*^2^ = 0.6). Moreover, the model separately estimated the magnitude of motor execution noise (0.11 ± 0.01 cm) and the cost of control (0.05 ± 0.02). Thus, our computational framework can jointly estimate the latent determinants of continuous sensorimotor behavior, isolating dynamic proprioception from other motor and cognitive processes.

### Dynamic and static proprioception constitute distinct somatosensory dimensions

It remains unclear whether dynamic proprioception and static position sense reflect common sensory capacity or dissociable somatosensory constructs (Lowrey et al., 2020; Proske, 2019; Proske C Gandevia, 2012; A. L. Wong et al., 2024; Zangakis et al., 2026). To address this question, a separate group of participants completed both our dynamic proprioceptive tracking task and a conventional static position-sense task (Experiment 2; N = 5). In the dynamic task, participants continuously tracked the unseen position of the right hand in both the mediolateral (x-axis) and anteroposterior (y-axis) dimensions (see Figure 4a for one participant’s cross-correlations in each dimension), with dynamic proprioceptive uncertainty estimated using our computational framework (Figure 3b). In the static task, participants completed a two-alternative forced-choice position discrimination task, judging whether the unseen right hand, relative to the unseen left hand, was farther from or closer to the body midline (x-axis) or farther from or closer to the body (y-axis). The function relating choice proportion to signed difference in position was used to estimate static proprioceptive uncertainty, as quantified by the just-noticeable difference (Straub C Rothkopf, 2022; Figure 4a).

**Figure 4.**
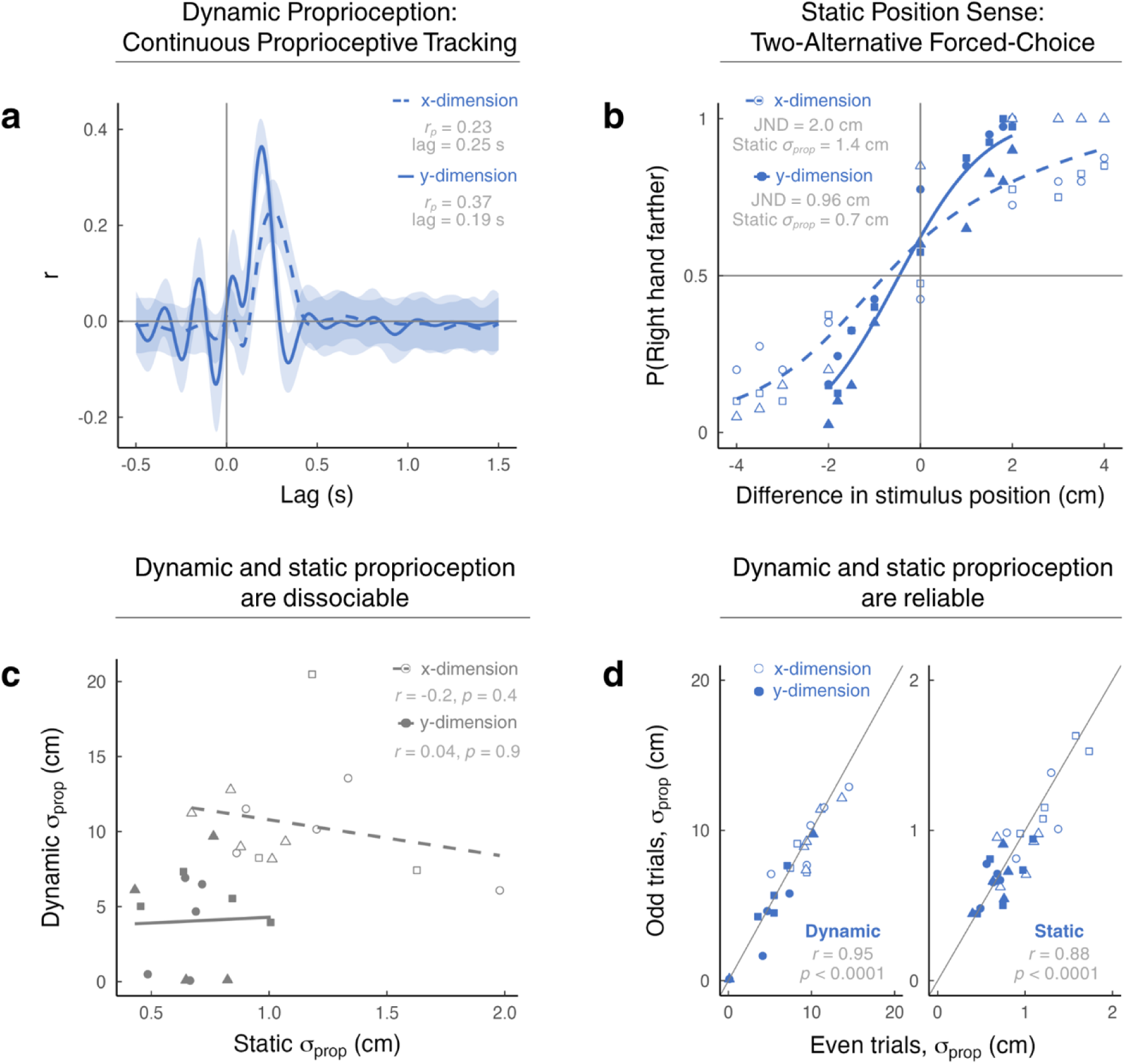
Dynamic and static proprioception constitute distinct somatosensory dimensions. (a-b) Representative data from a single participant. Left: Cross-correlation functions for the x-and y-dimensions during the dynamic tracking task; solid lines and shaded ribbons denote the mean ± SD. Right: Psychophysical function from the static position discrimination task, with the participant’s JND and corresponding static proprioceptive uncertainty (JND/√2) reported. **(c)** Dynamic and static proprioceptive uncertainty (*σ_prop_*) were not significantly correlated, indicating that the two assays quantify distinct dimensions of proprioceptive function. Regression lines are shown separately for each movement dimension, with solid lines denoting the y-dimension and dashed lines the x-dimension. **(d)** Split-half reliability of proprioceptive uncertainty for both assays, comparing estimates derived from odd-and even-numbered trials. Pearson correlation coefficients (r) and intraclass correlation coefficients (ICC) are reported. For b–d, points represent individual participants, with point shapes indicating workspace location; performance did not differ significantly across locations.

Importantly, the two tasks were designed to be as closely matched as possible, differing in whether proprioception was assessed during ongoing movement or at a static limb position. Across the two assays, this yielded 30 paired estimates of proprioceptive uncertainty (5 participants × 3 workspace locations × 2 movement dimensions). We varied movement dimension (x vs. y) to test whether the well-established anisotropy in static proprioception—poorer precision along the mediolateral (x) than the anteroposterior (y) axis—also characterizes dynamic proprioception. We also varied workspace location by shifting the tested region along the mediolateral and anteroposterior axes, allowing us to test whether proprioceptive precision—and the relationship between static and dynamic proprioception—generalizes across different regions of the workspace. Together, this design provided multiple paired estimates spanning movement dimensions and workspace locations with which to assess the relationship between static and dynamic proprioception.

We first asked whether the two assays captured common spatial features of proprioceptive processing. Across both tasks, proprioceptive uncertainty was lower along the anteroposterior (y) axis than the mediolateral (x) axis (*F*(1,42) = 23.9, *p* < 0.0001, *η*^2^ = 0.4; static: Δ = 0.5 ± 0.1 cm; dynamic: Δ = 5.7 ± 1.2 cm), consistent with previous reports of anisotropy in human proprioception (van Beers et al., 1998; Figure 4). This shared anisotropy suggests that directional differences in uncertainty reflect a fundamental property of proprioceptive processing rather than a feature specific to either assay. In contrast, workspace location had no reliable effect on proprioceptive uncertainty by either measure (*F*(2,42) = 0.9, *p* = 0.42).

Despite this shared spatial structure, dynamic and static proprioceptive uncertainty were not reliably associated in either the x-dimension (*r* =-0.2, 95% CI [-0.7, 0.4], *p* = 0.435) or the y-dimension (*r* = 0.04, 95% CI [-0.5, 0.6], *p* = 0.90; Figure 4c). Thus, individuals with more precise static position sense were not necessarily those with more precise dynamic proprioception. This dissociation should be interpreted cautiously given the modest sample size. Larger studies will therefore be important for establishing the robustness and generalizability of this dissociation.

Nevertheless, several features of the data argue against the lack of association arising simply from poor measurement. Dynamic proprioceptive uncertainty was estimated with high reliability (split-half reliability, r = 0.95; ICC(A,1) = 0.91, 95% CI [0.81-0.96]), as was static proprioceptive uncertainty (split-half reliability, r = 0.88, ICC(A,1) = 0.88, 95% CI [0.77–0.94]; Figure 4c). Neither measure showed evidence of floor or ceiling effects (dynamic proprioceptive uncertainty: 0.1–20.5 cm; static proprioceptive uncertainty: 0.4–2.0 cm).

Interestingly, dynamic proprioceptive uncertainty was significantly greater than static proprioceptive uncertainty (*F*(1,42) = 82.1, *p* < 0.0001, *η*^2^ = 0.66; estimated marginal means, dynamic vs static: 7.4 vs. 0.9 cm; *t*(38) = 9.1, *p* < 0.0001, *D_z_* = 1.4). One possibility is that continuously estimating limb state during ongoing movement imposes temporal demands that trade precision for speed. Together, the divergent individual estimates and overall levels of uncertainty suggest that dynamic proprioception and static position sense capture distinct dimensions of proprioceptive function.

## Discussion

Every movement unfolds with a simple question: Where is my body? The nervous system answers through proprioception—the sense of limb position at a given point in time (static position sense) and across continuous movement (dynamic proprioception). Although position sense has been well characterized, dynamic proprioception has remained difficult to measure directly. To overcome this limitation, we introduce a continuous tracking paradigm that, combined with computational modeling, provides a sensitive, reliable, and efficient assay of dynamic proprioception. Using this approach, we show that dynamic proprioception provides faster and more faithful estimates of limb state than vision and dominates multisensory state estimation during movement. Together, these findings establish a new paradigm for the study of proprioception in the context of ongoing movement, opening new avenues for understanding its role in health and disease.

An important feature of our approach is that proprioceptive state estimates are expressed continuously through behavior, without requiring explicit perceptual reports. The minimal sensorimotor lags observed during tracking further suggest that these estimates unfold rapidly and largely automatically. The resulting measures of individuals’ dynamic proprioceptive uncertainty were not reliably correlated with traditional measures of their static position sense based on explicit judgments of relative limb position, suggesting that the two assays may capture distinct dimensions of somatosensory function (Carranza et al., 2026; Chapman et al., 2001; Goble et al., 2012; Jones et al., 2012; McCloskey, 1973; Proske, 2025; Proske & Gandevia, 2009; Yasuda et al., 2014).

Our results also provide converging evidence that proprioception plays a privileged role in the dynamic state estimates underlying motor control (Cameron et al., 2014; Crevecoeur et al., 2016; Kasuga et al., 2022). During movement, dynamic proprioception provides faster and more precise estimates of limb state than vision, consistent with converging evidence showing that proprioceptive signals reach the motor system more rapidly than visual signals, that loss of proprioception is disproportionately disruptive to movement, and that corrective responses to mechanical perturbations are substantially faster than those to visual perturbations (Bair et al., 2002; Crevecoeur et al., 2016; Flanders et al., 1986; Ghez et al., 1995; Nowak et al., 1995; Pruszynski & Scott, 2012; Raiguel et al., 1989; Rothwell et al., 1982; Sainburg et al., 1993, 1995; Sanes et al., 1985; Sarlegna & Sainburg, 2009; Scott, 2012; Song & Francis, 2013).

This privileged role extended to multisensory state estimation. When visual and proprioceptive cues were aligned, as they typically are during natural movement, multisensory tracking closely resembled proprioception alone. Under cue conflict, vision exerted a smaller influence that scaled with its precision, consistent with reliability-weighted integration. Quantitative model comparison, however, could not clearly distinguish reliability weighting from near-exclusive reliance on proprioception. Future experiments that independently manipulate visual and proprioceptive reliability should provide greater leverage for resolving these possibilities (Eschelmuller et al., 2025; Grose et al., 2022). Additionally, more refined computational models incorporating realistic arm biomechanics, signal-dependent motor noise, and time-varying sensory delays or uncertainty may further sharpen this distinction.

Beyond these conceptual findings, continuous tracking offers practical advantages for quantifying proprioceptive function. Clinical examinations are often coarse, whereas laboratory psychophysical assays are typically time-intensive (Elangovan et al., 2014; Han et al., 2016; Horváth et al., 2023). Although robotic assessments have improved precision, their measurements frequently conflate proprioception with other motor and cognitive processes (Cressman et al., 2021; Kenzie et al., 2017; Park et al., 2023; Semrau et al., 2018; Tulimieri & Semrau, 2023). In contrast, our framework provides reliable estimates of dynamic proprioception in under several minutes and, through computational modeling, dissociates proprioceptive uncertainty from motor variability, control costs, and biomechanical constraints. The resulting assay is rapid, interpretable, and scalable, enabling quantitative assessment of dynamic proprioception across the lifespan and in neurological disease (Cao et al., 2025; Van De Plas & Orban de Xivry, 2026; Zimmet et al., 2020).

Several features of the current framework also define opportunities for further development. First, the current assay relies on bimanual matching, limiting its application to individuals who cannot use one limb to report the state of the other, including some patients with stroke and other motor impairments. Replacing manual matching with continuous eye tracking could extend the framework to these populations (Tulimieri et al., 2025). Second, the current assay probes proprioception primarily along two spatial dimensions (x and y), whereas natural movement unfolds across many degrees of freedom. Extending continuous tracking to complex, three-dimensional movements could reveal how dynamic proprioception operates during more naturalistic action. Third, we deliberately used unpredictable random-walk trajectories to minimize prediction and isolate feedback-driven proprioceptive estimation. Natural movements, however, are often predictable, structured, and learned (Ariani & Diedrichsen, 2019). Extending the paradigm to learnable sequences or self-generated movements could reveal how dynamic proprioception interacts with internal predictions during ongoing behavior (Aman et al., 2014).

## Conclusion

More than a century ago, Sherrington introduced proprioception as the sense of body position (static position sense) and movement (dynamic proprioception) (Sherrington, 1906). Yet while static position sense has been extensively characterized, far less is known about the rapid, continuous, and largely automatic computations that estimate the state of the body during movement. By combining continuous proprioceptive tracking with computational modeling, we make the latent dynamics of proprioception measurable, opening the door to understanding how the nervous system answers, from moment to moment, one of its most fundamental questions: Where is my body now?

## Methods

### Participants

Twenty participants took part in Experiment 1 (mean age = 20 years; all right-handed; 15 female), and six completed Experiment 2 (mean age = 24 years; all right-handed; 4 female). All had normal or corrected-to-normal vision and no history of neurological impairment. Participants provided written informed consent and received either course credit or $15 compensation. All procedures were approved by the Carnegie Mellon University Review Board (IRB: STUDY2024_00000279).

### Apparatus and General Procedure

Participants sat in front of a robotic manipulandum (KINARM endpoint 2D robot, BKIN Technologies, Kingston, Ontario, Canada) and grasped one handle with each hand. Movements were restricted to the horizontal plane and recorded at 1000 Hz. Visual stimuli were projected onto a horizontal mirror mounted above the workspace (refresh rate: 125 Hz), which occluded direct vision of the hands while creating the percept that visual stimuli and the hands occupied the same plane. Participants rested their forehead on a padded support throughout the experiment. Both experiments were conducted in darkness to eliminate external visual references. A Cartesian coordinate system was centered on the workspace, with positive x directed rightward and positive y directed away from the participant. Each trial began with participants positioning both handles within designated start locations (1.4-cm diameter circles) while receiving continuous visual feedback of hand position (1.4-cm diameter white circles). After both hands were maintained within the start locations for 1 s, the visual feedback and start markers disappeared and the trial began.

### Experiment 1 Procedure

In Experiment 1, the right hand always started at x = 12 cm, y = 17 cm, while the left-hand start position alternated randomly between two locations (x =-12 cm, y = 4 cm or x =-12 cm, y = 30 cm) to encourage active alignment between the hands.

Before each trial, participants were informed whether the target would be specified by proprioceptive or visual information. In the proprioceptive tracking condition, participants held both handles while the robotic manipulandum passively moved the right hand along a predefined random-walk trajectory. Participants continuously matched the vertical position of the right hand using their left hand (Figure 1a, first row).

In the visual tracking condition, participants released the right handle and rested their right hand by their side. A visual target (cursor or cloud) then appeared at the right-hand start location and moved along a predefined random-walk trajectory. Participants used their left hand to continuously track the target’s vertical position (Figure 1a, second and third row). Visual targets were rendered as two-dimensional Gaussian luminance blobs with either low uncertainty (cursor; *σ* = 30 pixels) or high uncertainty (cloud; *σ* = 180 pixels). To facilitate blending with the noisy mid-gray background, alpha masks were applied to each visual target, yielding peak opacities of 0.9 for the cursor and 0.4 for the cloud.

In the multisensory tracking condition, participants performed the same continuous tracking task while both a proprioceptive target (their right hand) and a visual target (either a cursor or a cloud) were presented simultaneously. Participants were told that the visual target represented the position of their right hand and were instructed to continuously track the position of their right hand.

For each trial, the target (either proprioceptive, visual, or multisensory) followed one of ten predefined trajectories generated as bounded one-dimensional random walks (for 3 example trajectories, see Figure 1b). Each trajectory was created by cumulatively summing 200 independent samples drawn from a zero-mean Gaussian distribution (SD = 1.2 cm). Trajectories were constrained to remain within y = 12–35 cm, with a maximum step-to-step displacement of 12 cm. The visual target’s trajectory was subsequently smoothed using a first-order low-pass filter to better approximate the continuous, naturalistic motion of the proprioceptive target. Position updates occurred every 77 ms (13.0 Hz), yielding a total trajectory duration of 15.4 s. At the end of each trial, the robot stopped moving and any visual stimuli disappeared. Trials were separated by a 20-s intertrial interval.

Participants completed two experimental sessions (∼40 min each) separated by an average of 7 days. Each session consisted of two phases: an aligned phase and an offset phase. During the aligned phase, participants completed 30 trials (10 proprioceptive, 10 visual, and 10 multisensory), presented in pseudorandom order such that each condition recurred every three trials. During the offset phase, participants first completed 26 multisensory trials in which the visual target was gradually displaced relative to the proprioceptive target by 0.5 cm per trial, reaching a final offset of 5 cm. They then completed an additional 30 trials (10 per tracking condition) with the 5 cm visuo-proprioceptive offset held constant. Across sessions, the proprioceptive condition was identical, whereas the visual and multisensory conditions differed only in the visual target: one session used a low-uncertainty visual cursor, and the other a high-uncertainty visual cloud.

### Experiment 2 Procedure

Experiment 2 consisted of two components: a dynamic proprioceptive assessment and a conventional static proprioceptive assessment. The dynamic proprioceptive assessment followed a similar procedure as the proprioceptive tracking condition in Experiment 1. Participants completed 120 proprioceptive tracking trials, with 40 trials in each of three workspace locations (Workspace 1 center: x = 12 cm, y = 12 cm; Workspace 2 center: x = 22 cm, y = 12 cm; Workspace 3 center: x = 12 cm, y = 22 cm; workspace extent: ± 10 cm about each center). The three workspace locations were tested in separate blocks and were chosen to sample distinct regions of the workspace. At the start of each block, both the left and right hands were positioned symmetrically about the center of the corresponding workspace. Participants were instructed to continuously mirror the position of their unseen right hand with their unseen left hand. Random-walk trajectories extended up to ±7 cm from the workspace center along both the x-and y-axes. Right-hand position updates occurred every 55 ms (18.2 Hz), yielding a total trajectory duration of 17 s.

The static proprioceptive assessment consisted of a two-alternative forced-choice (2AFC) position-discrimination task along the same three workspace locations as the dynamic assay (Wilson et al., 2010; J. D. Wong et al., 2011, 2014). Participants completed a total of 2160 trials in a single testing day (360 trials × 3 workspace locations × 2 axes; approximately 5 hours of testing). At the start of each trial, the KINARM robot passively moved both hands through four randomly selected positions over approximately 4 s to minimize positional cues. At the final position, participants judged whether their right hand, relative to their left hand, was farther from or closer to the body along the anteroposterior (y) axis, or farther from or closer to the body midline along the mediolateral (x) axis. Relative hand position was manipulated by systematically displacing the right hand with respect to the left by one of nine offsets (x-axis: ±4, ±3.5, ±3, ±2, 0 cm; y-axis: ±2, ±1.8, ±1.5, ±1, 0 cm), presented using the method of constant stimuli. Different offset ranges were used for the two axes because pilot testing revealed differences in proprioceptive sensitivity, allowing the sampled offsets to span the psychophysical function in each dimension (van Beers et al., 2002). Offset levels were presented in pseudorandom order, with each repeated once every nine trials. The order of workspace locations and of the dynamic and static assessments was counterbalanced across participants (see task videos, data, and code in our OpenMotor repository).

## Data analyses

### Participant-level exclusion

Participants were excluded if their tracking movements were not significantly correlated with the target trajectory, indicating failure to follow the tracking task instructions. In Experiment 1, one of 20 participants met this criterion, resulting in a final sample of 19 participants. In Experiment 2, one of six participants met the same criterion and was excluded, resulting in a final sample of five participants.

### Trial-level exclusion

In both experiments, the first 3 s of each trial were discarded to ensure analyses reflected steady-state tracking behavior. Then, dynamic proprioception trials were excluded if (1) participants loosened their grip on the right handle during proprioceptive trials, causing the robot to stop moving (Experiment 1: 0.17% of trials; Experiment 2: 0.08%); (2) the left hand remained stationary throughout the trial (Experiment 1: 0.09%; Experiment 2: 0); or (3) tracking performance was identified as an outlier (Experiment 1: 2.2%; Experiment 2: 4.1%). Outlier detection was performed separately for each experiment.

In Experiment 1, trials were grouped by participant, session, and condition; in Experiment 2, trials were grouped by participant, workspace location, and movement dimension. Within each grouping, trials with Fisher z-transformed peak cross-correlation values or corresponding cross-correlation lags exceeding ±2.5 standard deviations from the participant-specific mean were classified as outliers. Across all participants, 2.4% and 4.2% of trials were excluded in Experiment 1 and 2, respectively. In visual tracking trials, failure to release the right handle resulted in continued proprioceptive input, and these trials were therefore reclassified as multisensory trials (Experiment 1: 1.3%; not applicable to Experiment 2).

### Cross-correlation

The correspondence between target and hand movements was quantified using normalized cross-correlations between the velocity of the target and the left hand across temporal lags of ±1500 ms. For Experiment 1, cross-correlations were always computed relative to the visual target’s trajectory, which was displayed during visual trials and implemented as a hidden reference trajectory during proprioceptive trials that specified the robot’s motion; by establishing a common referent, this approach avoided subtle discrepancies between tracking modalities arising from the inertial dynamics of the robotic manipulandum. Correlation coefficients were Fisher z-transformed and averaged across trials at each lag, yielding a single cross-correlation function for each participant across modalities. For Experiment 2, cross-correlations were computed separately for each movement axis (x, y) and always between the left and right hands.

We extracted two key metrics from each averaged cross-correlation function (Figure 1c): First, the peak correlation coefficient, *r_p_*, quantified the strength of coupling between the left hand and the tracked target, with larger values indicating a stronger correspondence between the two signals. Second, the lag at the peak correlation quantified the temporal delay between target and tracking movements. Increased sensory uncertainty was expected to reduce the peak correlation and increase the lag, reflecting weaker coupling and longer processing delays, respectively (Burge & Bonnen, 2025).

### Reliability

In Experiment 1, the reliability of the cross-correlation metrics (peak correlation and lag) was quantified using two-way intraclass correlation coefficients for absolute agreement, single measures (ICC(A,1)). For proprioceptive tracking, reliability was assessed across sessions by comparing Session 1 and Session 2. For visual tracking, reliability was assessed within sessions by comparing estimates derived from even and odd trials. Pearson correlation coefficients (r) were additionally computed to facilitate visualization of the relationship between repeated measures. In Experiment 2, reliability was quantified using the same split-half approach. For dynamic proprioception, the computational model was refit separately to each half of the data to obtain independent estimates of proprioceptive uncertainty. For static position sense, psychometric functions were fit separately to each half, with proprioceptive uncertainty quantified as the just-noticeable difference (JND) divided by √2 (see below).

### Efficiency

We estimated the minimum number of trials required to obtain stable cross-correlation metrics in Experiment 1 using a bootstrap convergence analysis performed separately for each participant, session, and target modality. For each trial count (N = 1– 10), 1,000 bootstrap samples were generated by randomly sampling N trials with replacement. The mean estimate from each bootstrap sample was compared to the reference estimate obtained using all available trials. Convergence was defined as the smallest trial count for which the absolute relative deviation of the mean estimate from the reference estimate fell below 5% and remained below this threshold for all subsequent trial counts. Analyses were conducted separately for peak correlation and lag and only included proprioceptive and visual modalities.

### Tracking performance across modalities

In Experiment 1, peak correlation and lag for proprioceptive and visual tracking were analyzed using separate linear mixed-effects models (Kuznetsova et al., 2017). Fixed effects included the target modality (proprioceptive, visual cursor, visual cloud), and random effects included participant-specific intercepts. Model effects were evaluated using analysis of variance. Multisensory tracking error was analyzed using analogous mixed-effects models. To assess changes in proprioceptive tracking performance over the course of the aligned phase, trial-level root mean square error (RMSE) was calculated from the instantaneous tracking error within each trial. RMSE was analyzed using a linear mixed-effects model with trial number as a fixed effect and participant-specific intercepts as a random effect. Significant effects from the mixed-effects models were followed by two-tailed post hoc comparisons estimated with Tukey-adjusted p-values (Lenth et al., 2026). Effect sizes are reported as partial *η*^2^ for fixed effects, Cohen’s *D_z_* for within-subject pairwise comparisons, and Cohen’s *D* for one-sample comparisons against a reference value.

To characterize frequency-dependent attenuation of the proprioceptive tracking response, Fourier spectra were computed from passive right-hand and active left-hand trajectories using 10-s sliding windows after alignment using trial-specific lag at peak cross correlation. Active-to-passive gain was calculated across frequencies, and the slope of gain attenuation (20 log10[gain]) was estimated for each participant and tested against zero with a one-sample *t* test. Differences between multisensory cursor and cloud tracking during the offset phase were tested using a cluster-based permutation test (5,000 sign-flipping permutations), with candidate clusters defined as consecutive trials exceeding *p* < 0.05.

### Static vs. dynamic proprioceptive uncertainty

In Experiment 2, static proprioceptive uncertainty was estimated using a two-alternative forced-choice (2AFC) position-discrimination task. For each participant, workspace location, and axis, the proportion of responses indicating that the right hand was perceived as farther from the body (anteroposterior axis) or body midline (mediolateral axis) than the left hand was modeled as a function of the physical separation between the hands. A cumulative Gaussian psychometric function was fit using the quickpsy package (Linares & Lopez-Moliner, 2016). The just-noticeable-difference (JND) was then calculated from the psychometric curve as half the distance between the 75% and 25% response probabilities (JND = (*x*_75_ − *x*_25_)/2). Static proprioceptive uncertainty was then quantified as JND/√2, which estimates the uncertainty associated with a single position estimate (Kirsch & Kunde, 2019; Rohde et al., 2016; Straub & Rothkopf, 2022). Dynamic proprioceptive uncertainty (*σ_prop_*) was estimated from the computational model described below (see Description of the Computational Model). To assess the relationship between dynamic and static proprioceptive uncertainty, we fit a linear mixed-effects model with proprioceptive uncertainty estimates as the dependent variable, assay (dynamic vs. static), workspace location, and movement axis as fixed effects, and participant-specific intercepts as random effects.

### Description of Computational Models

Participants’ tracking behavior was modeled within an optimal feedback control framework using a linear–quadratic–Gaussian (LQG) controller (see Supplementary Materials for a full mathematical description). We considered four models that shared the same state-estimation, control, and biomechanical architecture but differed in how visual and proprioceptive information contributed to multisensory state estimation: proprioceptive-only, visual-only, equal-weighting, and reliability-weighting.

The environment was modeled as a discrete-time linear dynamical system in which the latent state evolved according to the previous state and motor commands, with additive Gaussian process noise. The system state included the target position (visual or proprioceptive), which evolved according to a Gaussian random walk imposed by the task design with fixed process noise (*σ_process_*), as well as the state of the tracking hand. At each time step, the agent received delayed, noisy observations of the target (visual, proprioceptive, or both) and the tracking hand (proprioceptive), corrupted by additive Gaussian sensory noise (free parameter, *σ*). Sensory delays were fixed at 150 ms for vision (Brenner & Smeets, 2003) and 75 ms for proprioception (Cameron et al., 2014; Kasuga et al., 2022).

The four models differed only in how visual and proprioceptive observations contributed to the target state estimate when both signals were available. The proprioceptive-only model relied exclusively on proprioceptive information, whereas the visual-only model relied exclusively on visual information. The equal-weighting model assigned equal weight to visual and proprioceptive observations regardless of their uncertainty. Finally, the reliability-weighted model weighted visual and proprioceptive observations according to their relative reliability, such that the more precise signal exerted greater influence on the resulting state estimate.

All four models shared the same control architecture. Actions were selected to minimize a quadratic cost function that balanced tracking accuracy (the difference between the target and tracking hand) against movement effort, with effort weighted by an action-cost parameter (*c*; free parameter) and motor commands corrupted by additive Gaussian execution noise (free parameter, *σ_motor_*). State estimation and control were implemented using a Kalman filter and linear quadratic regulator, respectively. The models additionally incorporated simplified arm biomechanics, in which motor commands generated forces that determined hand acceleration and velocity.

Models were fit to behavioral data using a Bayesian inverse modeling approach (Straub & Rothkopf, 2022). For Experiment 1, each model was fit jointly to the aligned phase from all sessions and experimental conditions for each participant, yielding a single set of parameters per participant. The model assumed the same control cost (*c*) and action variability (*σ_m_*_otor_) across conditions, reflecting the assumption that these parameters characterize stable properties of the participant rather than the sensory condition. The model included one proprioceptive uncertainty parameter (*σ_prop_*), and two visual uncertainty parameters (*σ*_cursor_, *σ*_cloud_) corresponding to the cursor and cloud visual targets. For Experiment 2, separate models were fit to the horizontal and vertical dimension and to odd and even trials to characterize reliability of the parameter estimates. Since there was no visual stimulus, the models for Experiment 2 had only three free parameters *(c, σ_motor_, σ_prop_)*.

Weakly informative priors were specified for all free parameters (see Supplemental Materials). Posterior inference was performed using the No-U-Turn Sampler (NUTS; Hoffman C Gelman, 2014) implemented in NumPyro (Phan et al., 2019), with four chains of 2,500 samples each following 2,000 warm-up iterations. Convergence was confirmed by R^<1.05 for all parameters (Gelman & Rubin, 1992). Model fits that failed to converge were excluded from the modeling analyses. This resulted in the exclusion of three participants in Experiment 1 and two data points in Experiment 2.

Expected log pointwise predictive density (ELPD) was used as a common measure of predictive accuracy across the two phases of Experiment 1. Because the models were fit to the aligned phase, aligned-phase ELPD was estimated using Pareto-smoothed importance sampling leave-one-out cross-validation (PSIS-LOO; Vehtari et al., 2017). The offset phase served as an independent held-out dataset, allowing ELPD to be computed directly from the log pointwise predictive density of the offset observations, averaged over posterior samples obtained from the aligned-phase fits. We compute differences in ELPD with respect to the reliability-weighting model, resulting in ΔELPD values for each participant. Overall model comparison was based on the mean and standard error of ΔELPD across participants.

## Data and code sharing

All data and analysis code associated with this study are publicly available on the OpenMotor repository (https://osf.io/aknqj/files/osfstorage).

## Acknowledgements

This work was supported by grant PID2023-150883NB-I00, funded by MICIU/AEI/10.13039/501100011033, to CM. PV was supported by grant PRE2021-097890, funded by MICIU/AEI/10.13039/501100011033 and the FSE+. JST was supported by National Science Foundation grant #2545300. We also thank Profs. Romeo Chua, Frédéric Crevecoeur, and Xaq Pitkow for their valuable advice during the early stages of this project.

---

**Figure S1.**
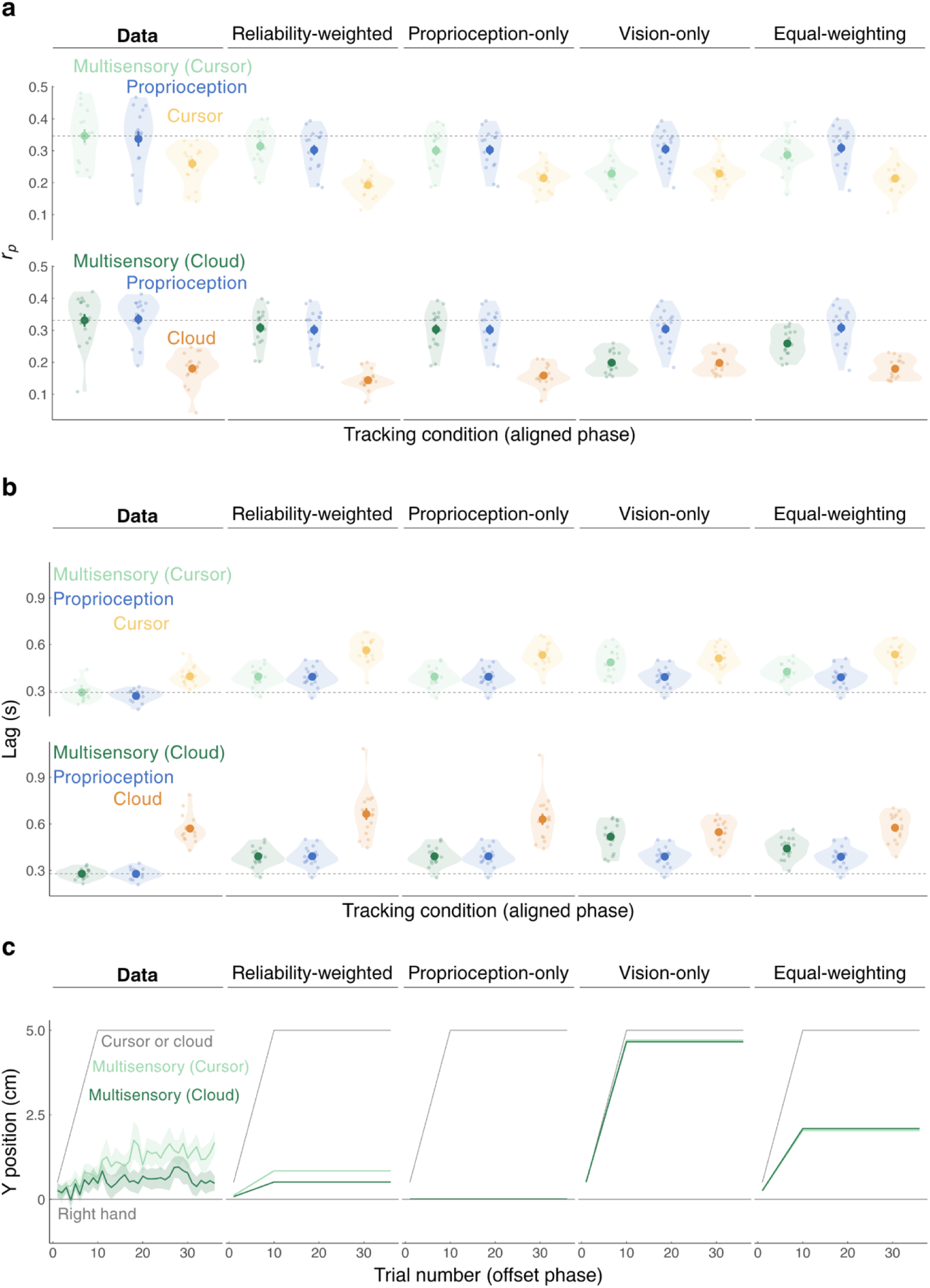
Posterior predictive checks for all four models. Model predictions for **(a)** cross-correlation and **(b)** lag during the aligned phase, and **(c)** tracking position during the offset phase. Translucent points show individual participant means; solid points and error bars show group means ± SEM. Model predictions are shown as lines and ribbons (mean ± SEM). Note that the vision-only model does not predict a full 5-cm shift toward the visual target, nor does the equal-weighting model predict a shift exactly to the 2.5-cm midpoint. These deviations arise from the shorter proprioceptive processing delay (75 ms) relative to vision (150 ms), which confers an inherent temporal advantage to proprioception during continuous tracking.

**Table S1.** Fitted model parameters. Parameter estimates for each of the four candidate models. Values represent group means ± SEM.

|  | Reliability-weighted | Proprioception-only | Vision-only | Equal-weighting |
| --- | --- | --- | --- | --- |
| $\sigma_{prop}$ | 4.3 ± 0.4 cm | 4.3 ± 0.4 cm | 4.2 ± 0.4 cm | 4.2 ± 0.4 cm |
| $\sigma_{cursor}$ | 8.2 ± 0.5 cm | 7.1 ± 0.4 cm | 6.4 ± 0.4 cm | 7.2 ± 0.5 cm |
| $\sigma_{cloud}$ | 12.2 ± 1.2 cm | 10.9 ± 1.1 cm | 7.7 ± 0.5 cm | 8.9 ± 0.6 cm |
| $\sigma_{motor}$ | 0.1 ± 0.01 cm | 0.1 ± 0.01 cm | 0.1 ± 0.01 cm | 0.1 ± 0.01 cm |
| **cost** | 0.05 ± 0.02 | 0.05 ± 0.02 | 0.05 ± 0.02 | 0.05 ± 0.02 |

## Supplementary material

## 1 Optimal control model details

### 1.1 General equations for LQG control

The state of the environment ***x****_t_* evolves according to a discrete-time linear dynamical system with additive Gaussian noise:

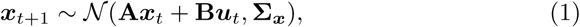

with noise covariance **Σ*_x_***.

At each time step, the agent receives a noisy observation ***y****_t_*of the state.

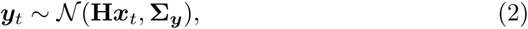

The observation is a linear transformation of the state plus additive Gaussian noise with covariance **Σ*_y_***.

The cost function is quadratic in the state and in the action:

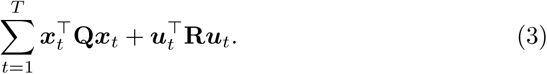

This control problem is known as linear-quadratic-Gaussian (LQG) control. Its optimal solution combines optimal state estimation via the Kalman filter (KF) (Kalman, 1960a) with optimal feedback control via the linear-quadratic regulator (LQR) (Kalman, 1960b).

The KF iteratively computes the posterior distribution over the state given observations, *p*(***x****_t_ |* ***y***_1_*,…,* ***y****_t_*) = *N* (***x̂****_t_,* **Σ***_t_*). The mean state estimate and its covariance are updated as

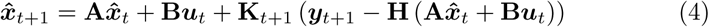

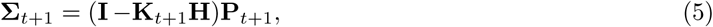

where **K***_t_* is

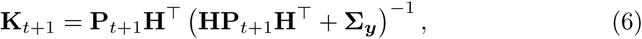

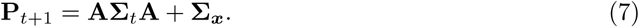

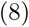

The linear-quadratic regulator yields a feedback policy expressed in terms of the current state estimate:

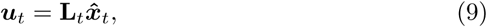

This feedback gain minimizes expected cumulative cost via dynamic programming; see e.g. Kochenderfer et al. (2022) for derivations of **L***_t_* and **K***_t_*.

### 1.2 Target and hand dynamics

We model the subject’s left hand as a point mass (position, velocity, and force states), while the target (right hand or visual target) follows a random walk with variance σ_rw_^2^. The model state is

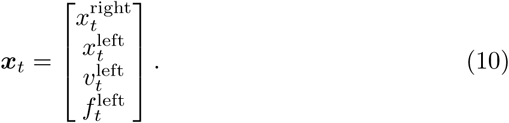

The left-hand dynamics are given by a point-mass model commonly used to describe reaching movements (see e.g. Kasuga et al., 2022, for details). Concretely, the continuous-time dynamics for the hand’s velocity and muscle force are

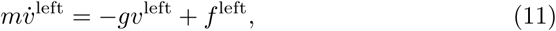

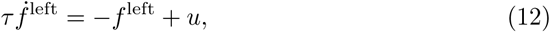

where *g* is a damping constant, *m* is the mass, *ε* is the muscle time constant, and *u* is the motor command. Together with *ẋ*^left^ = *v*^left^ these form a linear continuous-time system with matrices **A**_c_ and **B**_c_. Discretizing with time step *dt* (matrix exponential or zero-order hold) yields the discrete-time matrices **A**_pm_ and **B**_pm_. The continuous-time process noise is transformed to the discrete-time covariance **c**_pm_ via Van Loan’s method (Van Loan, 2003).

The target (right hand or visual target) is modeled as a Gaussian random walk with variance σ_rw_^2^:

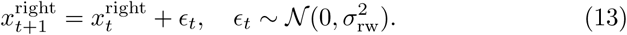

Stacking up the dynamics for the target and the left hand, the full discrete-time dynamics for the whole state are therefore

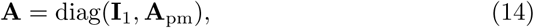

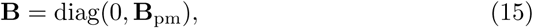

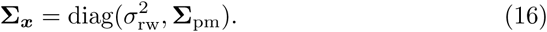

The cost function penalizes squared tracking error between response and target positions and squared control signals. The state and action cost matrices are

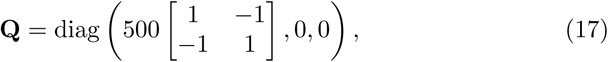

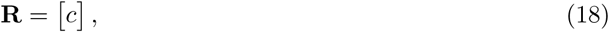

where *c* scales the action cost. The factor 500 in **Q** was chosen so that *c* remains in a numerically convenient range during fitting; since only the relative weighting of state and action costs matters, this does not change the model qualitatively.

### 1.3 Delayed multisensory observation model

We model sensory delays by augmenting the state vector (cf. Izawa & Shadmehr, 2008) and model multisensory observations as independent noisy channels for each sensory modality (cf. Crevecoeur et al., 2016; Kasuga et al., 2022).

Each sensory observation is the signed difference between right and left hand positions. Each modality *k ↓ {*prop, vis*}* has its own delay *ϖ_k_* and noise variance σ_k_^2^:

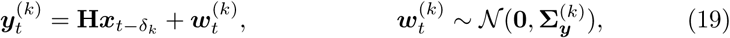

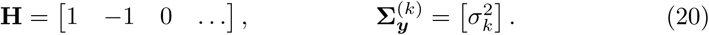

The complete sensory signal received by the agent is

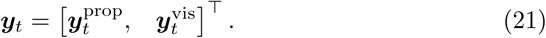

We set *δ*_vis_ = 2 and *δ*_prop_ = 1, which at our frame rate *dt* = 0.075 s corresponds to sensory delays of 150 ms (vision) and 75 ms (proprioception). The noise variances are free parameters and are estimated from the data.

To handle sensory delays, we augment the latent state by stacking previous states up to the maximum delay *δ*_max_ = max*_k_ δ_k_*and adjust the cost and dynamics matrices following Izawa and Shadmehr (2008) such that the previous *δ*_max_ states are stored in the state vector, but do not affect the dynamics or cost function:

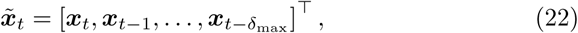

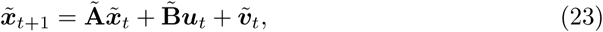

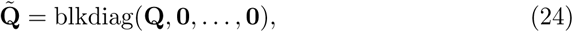

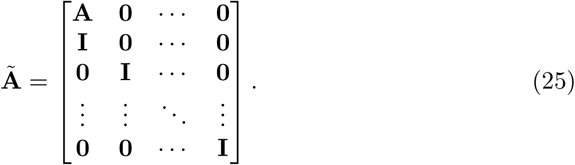

Each modality *k* then uses an augmented observation matrix H̃^(k)^ that selects the appropriately delayed component from ***x̃****_t_* (see Crevecoeur et al., 2016):

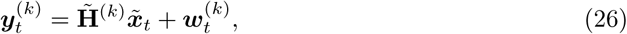

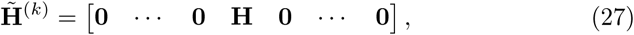

where the block **H** appears in the column block corresponding to delay *ϖ_k_*, and all other blocks are zero.

### 1.4 Model fitting

We fit the model to behavioral data using inverse optimal control (IOC) for LQG systems (Straub & Rothkopf, 2022). IOC defines the likelihood of a trajectory of observed states conditional on model parameters, 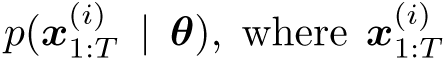 where denotes the *T*-step sequence of observed left-and right-hand positions in trial *i*. Individual trials are assumed independent and identically generated by the model with parameters ***θ***, so the dataset likelihood is 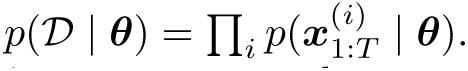.

To model the unisensory and multisensory conditions we assume the same action cost *c* and action variability *σ*_action_ across conditions. We also assume a single proprioceptive uncertainty parameter *σ*_prop_ and two visual uncertainty parameters, *σ*_vis,1_ and *σ*_vis,2_, one for each visual stimulus level.

Overall, the model has five free parameters

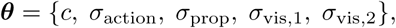

where *σ*_vis,1_ and *σ*_vis,2_ are the visual noise parameters for the two visual stimulus levels.

We use weakly informative priors for the parameters:

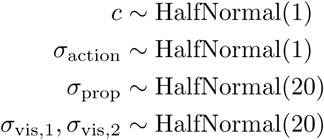

To test for optimal integration of visual and proprioceptive information we consider several model variants that differ only in how the multisensory condition is modeled; the unisensory conditions are modeled identically across variants.

1. In the *proprioceptive-only* model, only the proprioceptive sensory modality is used in the multisensory condition.
2. In the *visual-only* model, only the visual sensory modality is used in the multisensory condition.
3. In the *reliability-weighted integration* model, both visual and proprioceptive information is used in the multisensory condition and they are combined optimally as prescribed by a Kalman filter.
4. As a heuristic implementation of an *equal-integration* model, we assume that averaging happens at the level of parameters, such that both sensory modalities use the same parameter *σ_i_* = 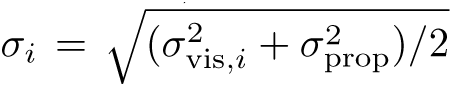 in the multisensory condition, while unisensory conditions still use their own modality-specific parameters *σ*_vis*,i*_ and *σ*_prop_, respectively.

### 1.5 Simulation and evaluation of the calibration phase

#### 1.5.1 Model with visual offset

To model a visual offset in the calibration phase of the experiment, we augmented the latent state with a scalar variable *b_t_* that represents the position of the visual cursor. This variable affects the visual observation but not the proprioceptive observation. If *x_t_*^base^ denotes the original latent state for either model, the augmented state is

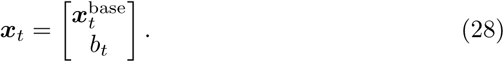

The proprioceptive observation matrix is

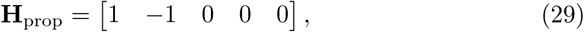

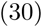

and the visual observation matrix is

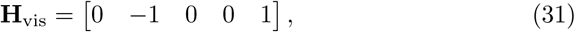

such that the visual element of the state affects only the visual observation. We assume that the agent is unaware of the bias, and the agent’s subjective internal model stays as in the original model without bias. The agent therefore does not expect a visual offset and does not maintain a separate belief about the visual cursor state.

#### 1.5.1 Simulation of the calibration phase

To simulate behavior in the calibration phase of the experiment, we took the 100 samples of the parameters from the posterior for each model and each participant. For each level of visual offset in the calibration phase, ranging from 0 to 5 cm in steps of 0.5 cm, we simulated one trial of tracking behavior per sampled parameter set. We averaged the tracking error (difference between left hand and visual target) across the 100 simulated trials to obtain the predicted tracking error for each level of visual offset.

#### 1.5.1 Evaluation of the calibration phase

To evaluate predictive performance, we evaluated the log-likelihood of the observed behavior in the calibration phase under the model with parameters sampled from the posterior (100 samples per model and participant).

